# Multi-dimensional diffusion MRI at ultra-high gradient strength for mapping axonal architecture and microstructure in the primate brain

**DOI:** 10.64898/2026.02.18.705310

**Authors:** Ting Gong, Chiara Maffei, Dongsuk Sung, Elissa Bell, Jingjing Wu, Jasmine Shao, Emma W Rosenblum, Xiangrui Zeng, Gabriel Ramos-Llorden, Alina Müller, Mirsad Mahmutovic, Boris Keil, Kabilar Gunalan, Satrajit Ghosh, Jean C Augustinack, Susie Y Huang, Suzanne N Haber, Anastasia Yendiki

**Affiliations:** Athinoula A. Martinos Center for Biomedical Imaging, Massachusetts General Hospital and Harvard Medical School, Charlestown, MA, United States; Mittelhessen University of Applied Sciences, Giessen, Germany; Massachusetts Institute of Technology, Cambridge, MA, United States; Department of Pharmacology and Physiology, University of Rochester, Rochester, NY, United States; McLean Hospital, Belmont, MA, United States

## Abstract

We present the most comprehensive sampling of the macaque and human brain with diffusion MRI to date. As part of the BRAIN CONNECTS center for Large-scale Imaging of Neural Circuits, we leverage ultra-high-gradient MRI systems, including the first-of-its-kind Connectome 2.0, for post-mortem acquisitions. Each sample is imaged for ∼250 hours at multiple spatial resolutions down to 0.25 mm isotropic for whole macaque brains and 0.4 mm isotropic for human hemispheres. Our optimized protocols allow us to sample both species across ∼50 diffusion shells varying in b-value, diffusion time, and echo time, reaching ultra-high b-values up to 64000 *s*/*mm^2^* with high signal-to-noise ratio. We demonstrate that these multi-dimensional data resolve not only white matter connectional architecture but also cortical and subcortical cytoarchitectonic boundaries, at a level of detail previously inaccessible in whole-brain noninvasive imaging. As such, these data are an important resource for both technical development and basic and clinical neuroscience.

## Introduction

Mapping primate brain circuitry at the whole-brain scale is essential for understanding the neural basis of cognition and behavior. Large-scale neural circuits comprise neurons in distant cortical and subcortical gray matter regions, interconnected by their axonal projections through the white matter. These connections underpin neural information processing, and disruptions to them have been linked to numerous neurological and psychiatric disorders^1–4^. Capturing both the cellular architectures of gray matter regions, and the organization of the white-matter fiber bundles connecting them, in the same, intact brains, is therefore fundamental not only for basic neuroscience but also for elucidating disease mechanisms and informing therapeutic interventions.

While there are microscopy techniques that can, in principle, achieve the sub-micron resolution needed to resolve and quantify axonal and cellular microstructure, such techniques do not scale readily to whole primate brains in terms of either acquisition or analysis. Even in limited fields of view, the need for tissue preparation, sectioning, and subsequent image analysis, including stitching, feature segmentation, and proofreading, introduce substantial burdens. As these microscopic techniques continue to be developed, access to whole-brain microstructural maps and wiring diagrams at a mesoscopic scale is essential to assist with localizing structures of interest and guiding image acquisition, analysis, and interpretation for microscopy.

Diffusion MRI (dMRI) provides a complementary, whole-brain and non-invasive approach for probing axonal and cellular architecture by sensitizing the MRI signal to the motion of water molecules within tissue microenvironments^5^. When combined with biophysical modeling, which disentangles the signal arising from different tissue compartments in a voxel, dMRI can probe microscopic features orders of magnitude smaller than the voxel size. These include compartmental tissue properties such as cell and neurite densities and geometries^6,7^, as well as fiber orientation distributions that can be used to reconstruct long-range connections via tractography^8^. Furthermore, whereas *in vivo* dMRI is constrained by acquisition time and subject motion, typically limiting resolution to several millimeters, post-mortem imaging enables ultra-long acquisitions that push spatial resolution into the sub-millimeter regime for deep phenotyping of primate brain organization^9^.

Despite these advantages, post-mortem dMRI faces technical challenges that limit the sensitivity of existing datasets, even the highest-resolution ones, to microstructural features. Tissue extraction and fixation markedly reduce T2 relaxation time and water diffusivity^10^, leading to competing demands for shorter echo times (TEs) and greater diffusion weightings (b-values), to ensure sensitivity to tissue microstructure while maintaining sufficient signal-to-noise ratio (SNR). These demands become particularly difficult to satisfy at high spatial resolutions. As a result, despite substantial efforts across species from marmosets to humans^11–15^, current post-mortem dMRI datasets in primate brains still lack the combination of ultra-high resolution and ultra-high b-values needed to resolve connectional and microstructural features that span the mesoscale-to-cellular range.

The ability of an MRI scanner to achieve such high b-values, while maintaining high spatial resolution, is limited by the maximum strength of its gradient field. Until recently, ultra-high gradient strengths were available only on small-bore animal MRI systems. With the development of the Connectome 2.0 system^16,17^, a 3T human MRI scanner equipped with 500 mT/m gradient strength, and similar systems^18^, it is now possible to achieve diffusion sensitivity similar to that of animal scanners across the whole human brain. In this work, we leverage these developments to perform the most comprehensive sampling of the human and macaque brain to date with dMRI. A multi-dimensional acquisition across multiple spatial resolutions is designed to reveal both axonal architecture and tissue microstructure in unprecedented detail, support systematic cross-species neuroanatomical investigations, and provide a testbed for further development of advanced dMRI modeling methods.

In the following, we show results from two human hemispheres and four macaque brains. These are the first specimens imaged for the BRAIN CONNECTS Center for Large-Scale Imaging of Neural Circuits, and will subsequently undergo optical and X-ray microscopy, serving as a unique, correlative imaging resource. We summarize the dMRI acquisition protocols and showcase representative analyses of connectional anatomy and microstructural features of various scales across the brain, demonstrating the image quality and potential of these datasets. Both the dMRI data and the derivative maps presented in this paper will be available at https://gallery.lincbrain.org.

## Results

### Acquisition protocol optimization

We imaged whole human hemispheres on the Siemens Connectome 2.0 3T MRI scanner with maximum gradient strength G_max_ = 500 mT/m and a custom-built *ex vivo* imaging coil^19^, and whole macaque brains on a Bruker small-bore 4.7T MRI system with maximum gradient strength G_max_ = 660 mT/m. For tractography, our goal was to achieve as high spatial resolution as possible, while reaching the high-b regime. For microstructural modeling, our goal was to achieve as high b-value with as many b-values, diffusion times, and echo times as possible. Rather than trading off between these requirements, we opted to develop separate protocols for these two goals. We refer to them as high-resolution and multi-dimensional protocols, respectively.

All dMRI data are acquired using a segmented 3D echo planar imaging (EPI) sequence^9^ with a high number of segments to minimize distortions and maximize SNR^20^. In addition, we acquire multiple spin-echo images for T2 mapping in each specimen. This allows us to evaluate the T2 relaxation time in gray and white matter for each species and thus optimize echo times used in the dMRI scans. This optimization aims at good tissue contrast and high SNR in all dMRI protocols, and sufficient sampling of the T2 decay curve in the multi-echo dMRI protocols for combined diffusion-relaxometry analysis. The resulting datasets span multiple spatial resolutions and diffusion-encoding dimensions in both species (Fig.1 and Extended Data Fig.1).

**Figure 1.**
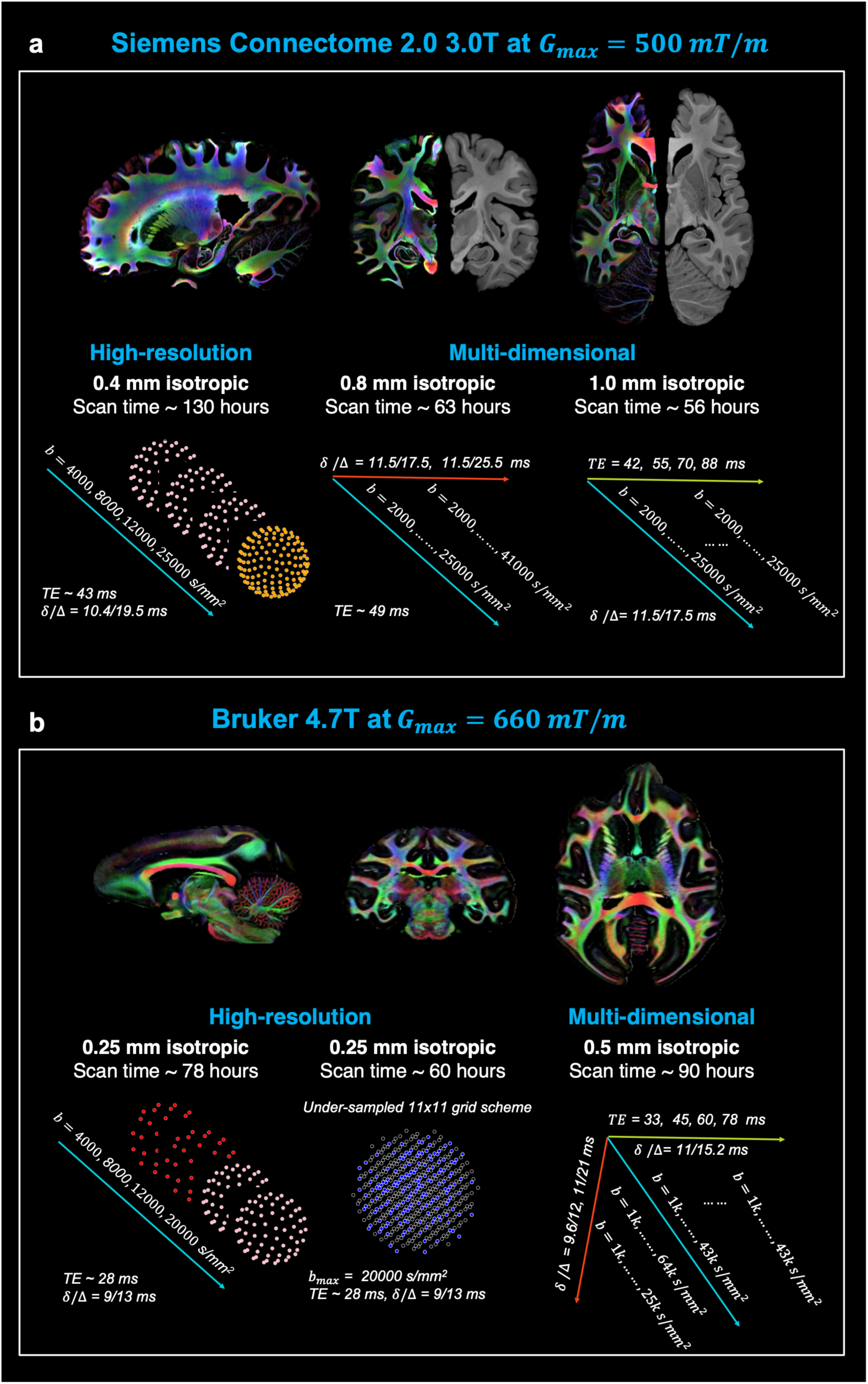
Summary of acquisition protocols for human and macaque dMRI. Direction-encoded color maps derived from the high-resolution data are shown for demonstration. **(a) Human hemispheres: i)** The high-resolution protocol features 0.4 mm isotropic resolution with four b-shells sampled at 64, 64, 64, and 128 diffusion directions, respectively. **ii)** The multi-dimensional protocols include two separate acquisitions: a two-diffusion-time scan (six b-shells at shorter diffusion times and eight b-shells at longer diffusion times, each with 32 or 64 directions) and a four-echo-time scan (six b-shells per TE, each with 32 or 64 directions). **(b) Macaque brains: i)** The high-resolution protocols feature 0.25 mm isotropic resolution with four b-shells sampled at 32, 32, 64, and 64 diffusion directions, respectively. In addition, high-resolution data are acquired with 171 directions using a three-times under-sampled Cartesian grid scheme to recover full diffusion spectrum data. **ii)** The multi-dimensional protocol included multiple diffusion times and echo times in a single scan. Detailed acquisition parameters including T2 values of the specimens, are provided in Extended Fig.1.

#### High-resolution protocols

These protocols prioritized spatial resolution, while maintaining sufficient SNR and angular resolution, to resolve small and deep fiber bundles. For human hemispheres, we achieved 0.4 mm isotropic resolution with a maximum b-value of 25000 *s*/*mm*^2^ and a short TE of 43.0 ms, at an average temporal SNR of ∼ 25. For macaque brains, we achieved 0.25 mm isotropic resolution with a maximum b-value of 20000 *s*/*mm*^2^ and a short TE of 28.4 ms, yielding an average SNR of ∼45. Data from both species contain at least four spherical shells with different b-values.

#### Multi-dimensional protocols

These protocols sampled the diffusion signal not only across gradient directions and b-values, but also across diffusion times and TEs, critical for advanced biophysical modeling, such as combined diffusion-relaxometry analysis^21^. We relaxed the demands on spatial resolution slightly to allocate scan time toward improving SNR at ultra-high b-values and longer TEs.

For human samples, we divide the multi-dimensional sampling in two due to scan time constraints: a multi-diffusion-time protocol at 0.8 mm isotropic resolution (6 b-shells at *δ*/Δ = 11.5/17.5 ms and maximum b-value of 25000 *s*/*mm^2^*; 8 b-shells at *δ*/Δ = 11.5/25.5 ms and maximum b-value of 41000 *s*/*mm^2^*; both at TE = 49 ms) and a multi-TE protocol at 1.0 mm isotropic resolution (four TEs at 42, 55, 70 and 88 ms, each with 6 b-shells with maximum b-value 25000 *s*/*mm^2^*, *δ*/Δ = 11.5/17.5 ms).

For macaque brains, the smaller sample size allows us to optimize a single, multi-diffusion-time and multi-TE protocol at 0.5 mm isotropic resolution. The protocol includes six consecutive acquisitions: TE = 28 ms, *δ*/Δ = 9.6/12.0 ms, maximum b-value of 25000 *s*/*mm^2^*; TE = 33 ms, *δ*/Δ = 11.0/15.2 ms, maximum b-value of 43000 *s*/*mm^2^*; TE = 38 ms, *δ*/Δ = 11.0/21 ms, maximum b-value of 64000 *s*/*mm^2^*; and three sessions at TE = 45, 60, 78 ms and *δ*/Δ = 11.0/15.2 ms with maximum b-value of 43000 *s*/*mm^2^*. These protocols are designed to utilize the maximum gradient strength (G_max_) to reach the highest *b*-values at the shortest possible diffusion times and, consequently, the shortest achievable TEs. Lower maximum *b*-values correspond to shorter TEs.

Summary of SNR estimates in each dataset is included in Extended Data Table 1-2.

### High-resolution tractography resolves detailed axonal architecture

The high-resolution datasets captured fiber organization in great detail, even in complex regions such as the thalamus, hippocampus, cerebellum, and brainstem in both species, as visualized by the direction-encoded color maps^22^ from fiber orientation distributions (Fig. 2a-b). The data also revealed clear transitions of major fiber orientations at the gray-white matter interface (Extended Data Fig.2), which are critical for reconstructing superficial white-matter tracts.

**Figure 2.**
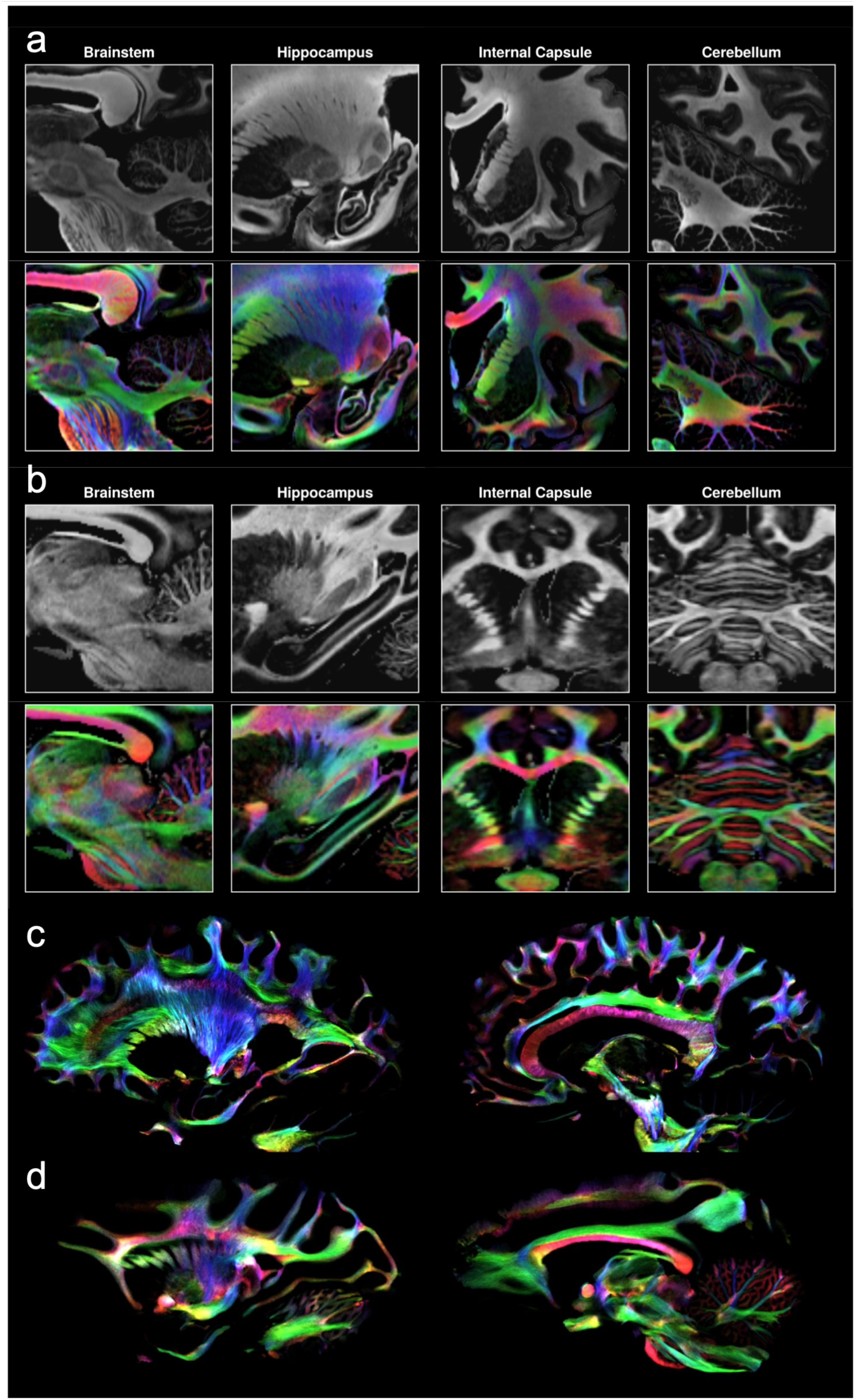
Detailed axonal architecture revealed by the high-resolution data in human (a, c) and macaque (b, d). The isotropic component (in gray contrast) and direction-encoded color maps (in RGB) from fiber orientation distribution functions (fODFs) in complex regions in human **(a)** and macaque **(b);** tract density images derived from whole-brain probabilistic tractography in human **(c)** and macaque **(d)**. The images are color-coded by fiber orientation (red for left-right, green for anterior-posterior, and blue for superior-inferior).

The tract density images derived from whole-brain probabilistic tractography further showed detailed axonal architecture across the white-matter, demonstrating the ultra-high spatial resolution of the data in both human and macaque (Fig.2c-d). We segmented specific fiber tracts to demonstrate complex topographical rules of smaller fiber bundles from different cortical areas as they travel within the same white-matter pathways, replicating results from invasive anatomic studies.

#### Cortico-subcortical projections through the internal capsule

High-resolution tractography delineated prefrontal cortex projections to the thalamus and brainstem via the internal capsule (Fig.3a-b), a densely packed white-matter structure. Tracts originating from distinct prefrontal regions were clearly separated as they travel within the capsule, which is not possible at conventional dMRI resolutions. Furthermore, the relative position of the fibers within the capsule aligned with known topographic rules from anatomic tracer studies in macaques^23^.

**Figure 3.**
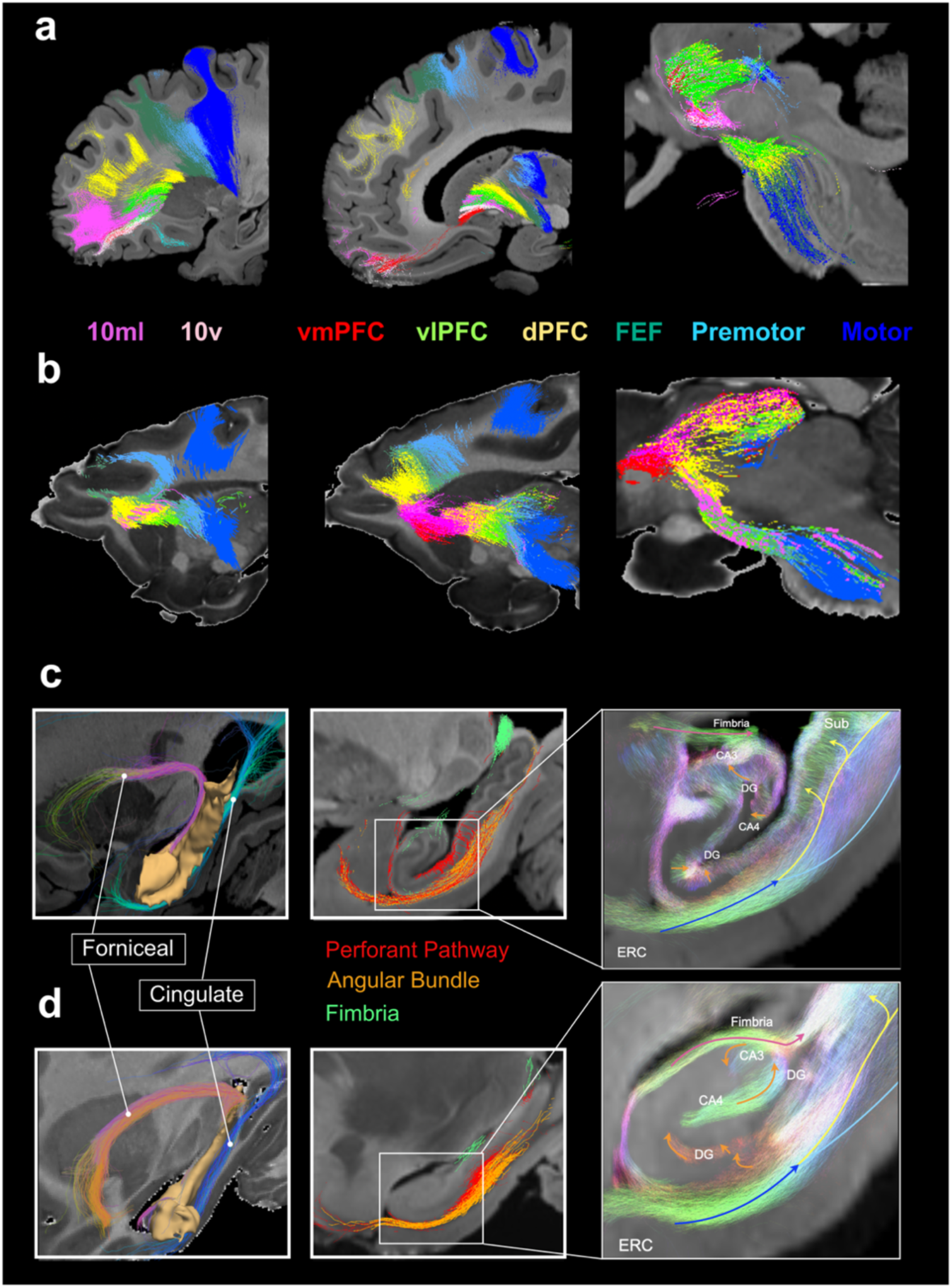
White-matter architecture in human (a, c) and macaque (b, d). Cortico-subcortical connections that travel through the internal capsule are shown in human **(a)** and macaque **(b),** at three levels: as fibers leave the prefrontal cortex (left), enter the internal capsule (middle), and leave the capsule to enter the brainstem (right). Tracts are colored according to their cortical terminations: medial and lateral area 10 (10ml), ventral area 10 (10v), ventromedial prefrontal cortex (vmPFC), ventrolateral PFC (vlPFC), dorsal PFC (dPFC), and frontal eye field (FEF). Hippocampal tracts of multiple spatial scales are shown in human **(c)** and macaque **(d)**, from the larger forniceal and cingulate bundles (left), to the intermediate-size hippocampal pathways (middle), to the small, intra-hippocampal mossy fibers and Schaffer collaterals (right; see also Extended Data Fig.3). Regions of interest include the entorhinal cortex (ERC), dentate gyrus (DG) and cornu ammonis (CA) fields.

#### Hippocampal tracts

With more densely seeded tractography within the hippocampus, we could resolve progressively deeper and finer connections to and within the hippocampus (Fig.3c-d). In both human and macaque brains, the high-resolution data enabled visualization of long-range and intra-hippocampal pathways spanning multiple spatial scales. These ranged from large forniceal bundles (originating primarily from CA1 and subiculum and projecting to the fornix) and cingulate bundles (entorhinal cortex to cingulum), to intermediate hippocampal pathways, including the perforant pathway, angular bundle, and fimbria, and to fine intra-hippocampal connections, including mossy fibers and Schaffer collaterals (Extended Data Fig.3). The organization of these pathways is consistent with hippocampal anatomy observed in microscopy^24^.

### High-resolution microstructure mapping delineates cytoarchitectonic layers

The human high-resolution data had a higher effective resolution than the macaque data when accounting for differences in brain size and cortical thickness, enabling delineation of cytoarchitectonic layers in both cortical and subcortical structures. To demonstrate this capability, we fitted the Soma and Neurite Density Imaging (SANDI) model^25^, adapted to *ex vivo* dMRI with the addition of a “dot” compartment^20^, to visualize cytoarchitectonic boundaries. This analysis allowed the estimation of the intra-soma and intra-neurite signal fraction, but requires multiple, high b-values, which can only be acquired with ultra-high gradient strength. We have implemented a probabilistic neural network estimator in the Microstructure.jl package, which allows us to fit this model to large, high-resolution datasets in a computationally efficient manner^26^.

#### Hippocampal subfields and substructures

The high-resolution SANDI maps allowed us to differentiate between hippocampal substructures (Fig.4). In the dentate gyrus, the granular layer exhibited high intra-soma signal fraction, while the surrounding stratum radiatum/stratum lacunosum-moleculare (SRLM) showed low intra-soma but high intra-neurite signal fraction, consistent with the cell packing density revealed in histology^27^. The alveus, a thin sheet of white matter covering the ventricular surface of the hippocampus, and the fiber-rich fimbria, were also visible in the intra-neurite signal fraction map. Distributions of microstructural measures within each segmented subfield further support our observations quantitatively. Specifically, CA4 showed a more centered distribution of intra-soma signal fraction with narrower interquartile range compared to CA1-CA3. This agreed with histology findings showing that CA4 has more evenly distributed neurons, while CA1-CA3 have greater variations of cell densities across layers. Fig.4 shows results from human specimen Ha1. While Hb1 exhibited a wider distribution of both measures in each region (Extended Data Fig.4), likely due to lower SNR compared to Ha1, the relative relationship among regions was the same.

**Figure 4.**
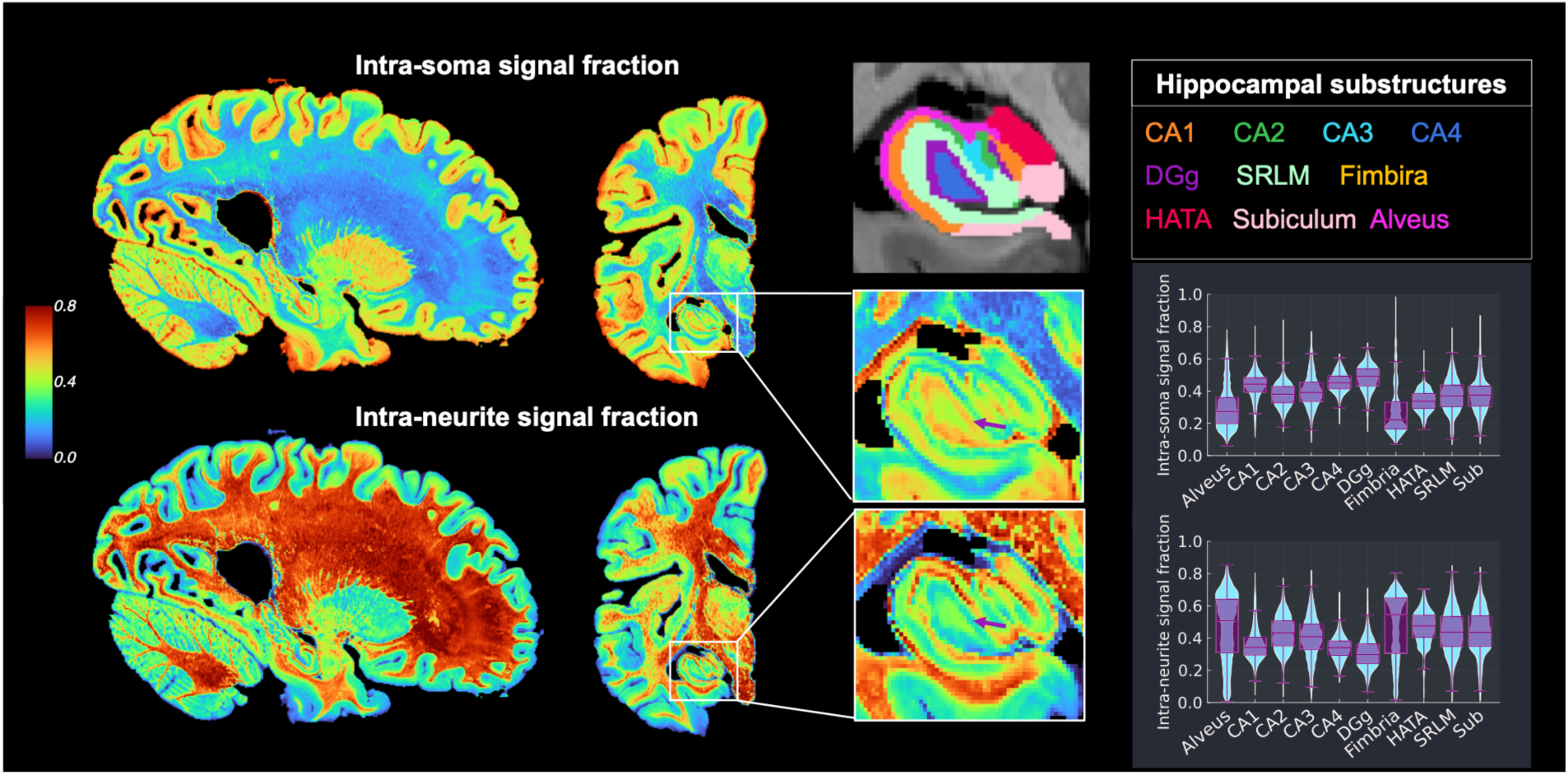
Gray-matter microstructure variation within the hippocampus. The intra-soma signal fraction and intra-neurite signal fraction derived from the high-resolution human data reveals cytoarchitectonic variations across hippocampal substructures. Manually annotated subregions of the hippocampus include cornu ammonis field 1-4 (CA1-CA4), subiculum, hippocampal-amygdaloid transition area (HATA), alveus, fimbria, granular layer of dentate gyrus (DGg) and the stratum radiatum and stratum lacunosum-moleculare (SRLM). The arrows in zoom-in view highlight the granular layer DGg with dense cell packing. Distributions of microstructure parameters from all voxels within each annotated region of the hippocampus are given as violin and boxplots (right; see also Extended Data Figure 4).

#### Cerebral cortex

The cerebral cortex has six cytoarchitectonic layers (I-VI). With ultra-high-resolution *ex vivo* structural images (0.12 mm isotropic resolution at 7T), a previous study was able to resolve the boundary between supragranular (layer I-VI) and infragranular (layer V-VI) layers^28^, showing quantitative agreement with segmentations from histology in the BigBrain dataset^29^. This boundary is also visible in our 0.4 mm isotropic T2-weighted mean b=0 image. We applied a semi-supervised deep-learning approach^28^, previously developed for structural images of the human brain, to identify this boundary in the mean b=0 images of our high-resolution human dMRI data, after up-sampling them to 0.12 mm isotropic resolution (Fig.5). Our maps show further cytoarchitectonic variations within the segmented supragranular and infragranular layers, demonstrating sensitivity to microstructure that is not present in standard structural images even with higher spatial resolution.

**Figure 5.**
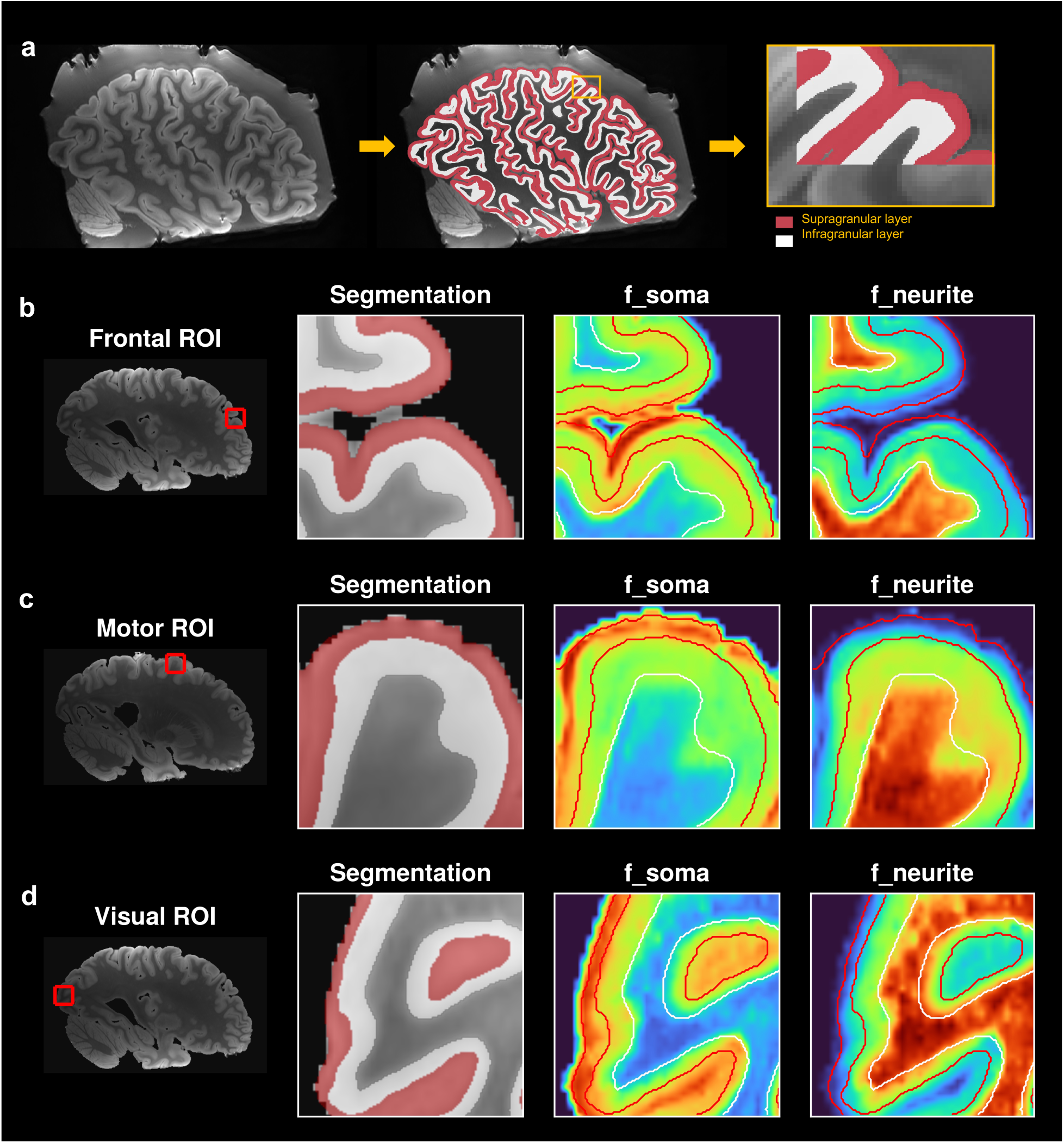
Gray-matter microstructure variation within cortical layers. **(a)** Segmentation of the supragranular and infragranular layer of the cerebral cortex based on the mean b=0 images from the high-resolution human data. **(b-d)** Zoom-in regions showing microstructure variations within segmented layers in the prefrontal cortex, motor cortex, and visual cortex.

Layer I of the primate cortex contains very few neuronal cell bodies and is extremely thin, typically less than 0.4 mm^29^. Layer II, although also thin, in general exhibits high cell-body density varying across the cortex. In our data, Layer II was well represented in the intra-soma signal fraction, appearing as a prominent high-signal band located immediately below the outer boundary of the segmented supragranular layer. Layer I is generally difficult to visualize directly at 0.4 mm isotropic resolution. Nevertheless, its influence can still be detected in some cortical regions as a subtle, slightly lower-signal band at the outer boundary of the supragranular layer. This is expected due to partial-volume effects between a very thin molecular layer and the immediately adjacent Layer II.

Comparing these microstructural features between cortical regions, we observed a relatively lower intra-soma signal fraction band below the high-signal strip in the primary motor cortex, whereas primary visual cortex showed a more uniformly high supragranular intra-soma signal fraction. This pattern could correspond to two histological findings: (i) primary visual cortex has a cell-dense Layer IV that contributes to a consolidated high cell density profile^30^, while the primary motor cortex is functionally agranular, lacking layer IV and (ii) the primary motor cortex is characterized by lower overall neuron packing density and relatively higher intracortical myelin content, factors that reduce the relative contribution of soma signal. In comparison, prefrontal cortex showed a heterogeneous but overall lower intra-soma fraction in the supragranular layers in our data, which agrees with histological studies^31^.

Finally, infragranular layers (V–VI) tended to exhibit lower intra-soma fraction than supragranular layers in visual and especially motor cortex in our measurements. This is consistent with histology showing that (i) the densest laminar compartment of the visual cortex is Layer IV (above Layer V), and (ii) motor cortex has hypertrophied Layer V with large pyramidal cells but overall lower neuron packing density in infragranular layers relative to the peak granular densities of sensory cortex. In prefrontal cortex, microstructural differences between the infragranular and supragranular layers were smaller and more variable across areas.

### Multi-dimensional data advance microstructural sensitivity and specificity

Complementary to the high-resolution and high-b-value data, sampling of the dMRI signal across the TE and diffusion time dimensions reduces bias in compartment volume fraction estimates, thus improving sensitivity to tissue composition and compartment size. Our comprehensive multi-dimensional protocol allows the application of a variety of previously proposed biophysical models and may enable the evaluation of new models. Here, we show some representative analyses.

#### Disentangling tissue microstructure and composition

Fig.6(a-b) compares microstructure maps estimated by fitting multi-TE SANDI (MTE-SANDI)^32^ vs. conventional, single-TE SANDI in the macaque. The latter is known to yield biased (T2-weighted) compartment signal fractions when T2 differences exist between compartments. We used data from the macaque multi-dimensional protocol with TE = 33, 45, 60, and 78 ms and same diffusion times, and performed parameter estimation using Microstructure.jl^26^.

**Figure 6.**
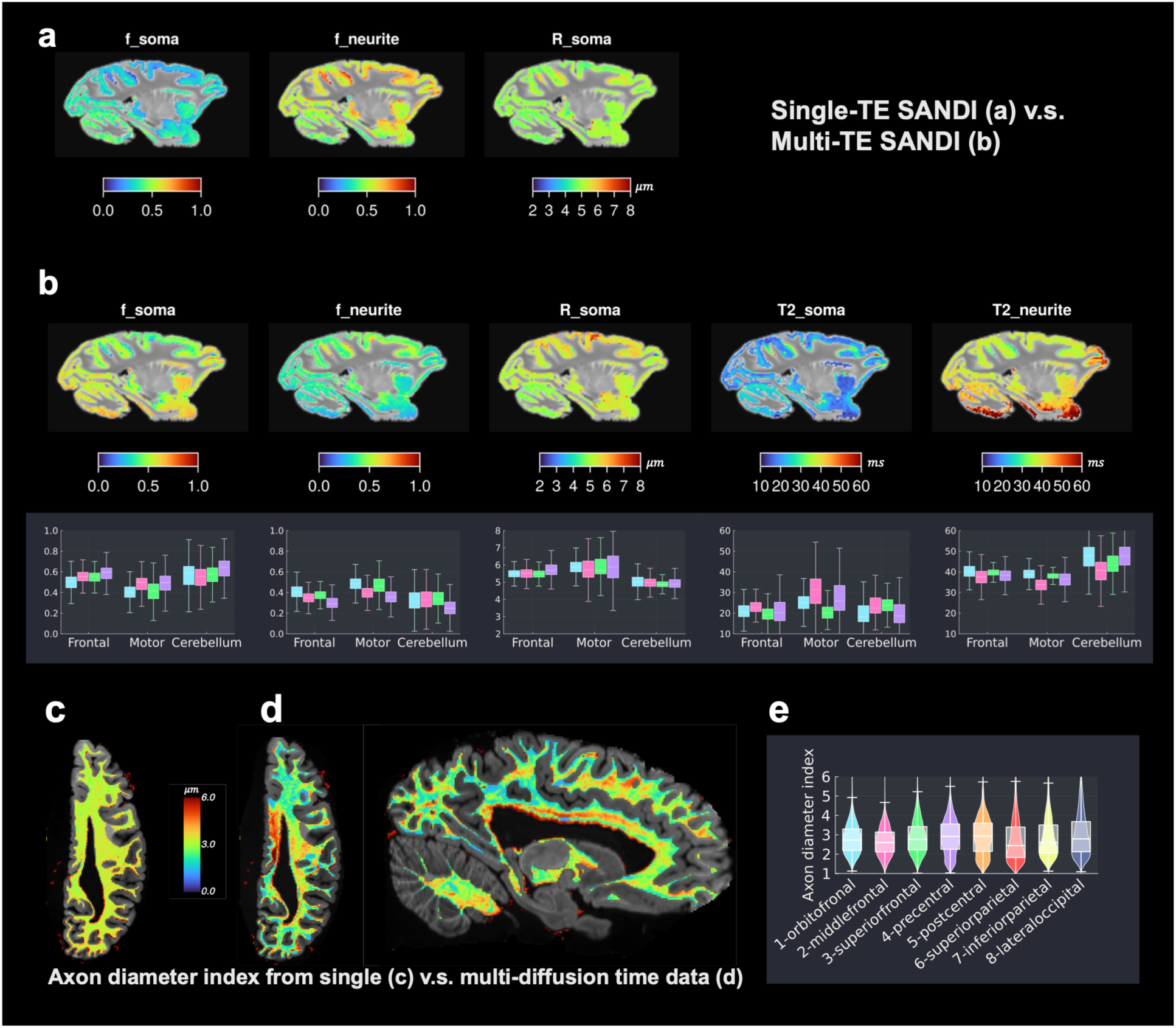
Gray- and white-matter microstructure imaging with multi-dimensional dMRI data. (a-b) Microstructural parameters in the GM, derived by fitting the SANDI model to conventional single-TE (a) vs. multi-TE (b) macaque dMRI data. The single-TE SANDI model derives T2-weighted intra-soma signal fraction (f_soma), intra-neurite signal fraction (f_neurite) and soma radius (R_soma); MTE-SANDI derives more accurate intra-soma and intra-neurite fractions, while also estimating the T2 values of the soma and neurite compartments. Boxplots from four macaques (b) show consistent variabilities in ROIs of the frontal cortex, motor cortex and cerebellum gray matter. (c-e) Axon diameter index in the white matter in the human brain with single-diffusion time dMRI data (c) and two diffusion-time dMRI data (d), and the variation of the latter across white-matter regions from the frontal to the occipital white-matter (e).

The soma compartment exhibited lower T2 values in gray-matter compared to the neurite compartment, consistent with previous observations from an *in vivo* human study^32^. This resulted in under-estimated intra-soma volume fraction and overestimated intra-neurite volume fraction, the level of which depends on the T2 differences between compartments. Notably, the non-T2 weighted (and therefore less biased) soma volume fraction map obtained with MTE-SANDI highlighted the cerebellum, reflecting its densely packed cells and smaller cell bodies. The soma radius map from MTE-SANDI also detected larger cell bodies in the motor cortex.

#### Mapping axon diameter index

Fig.6 shows the axon diameter index estimated from the two-diffusion time data in the human brain, using a three-compartment model of the white-matter^20^, including a cylinder compartment representing axons, a zeppelin compartment for extra-cellular space, and the dot compartment accounting for immobile water, typically present in the white-matter *ex vivo*. We used a Markov Chain Monte Carlo (MCMC) sampler for parameter estimation^26^. Prior studies have shown that estimating this microstructural parameter is challenging, and even the highest gradient strengths available in today’s high-performance MRI scanners are not sufficient for accessing the full range of axon diameters in the primate brain^33,34^. Hence these models do not aim to estimate the absolute value of axon calibers, but rather a relative index capturing differences between brain regions and individuals.

Encouragingly, the axon diameter index maps showed the sensitivity of the multi-diffusion-time protocol to differences in the axon diameter index across different brain regions and set the stage for further research on this topic. For example, while human sample Hb1 have sufficient SNR to support voxel-wise fitting of the axon diameter index, an alternative approach is to fit this parameter to signals averaged over tract segments^35^, which may further improve the robustness of the spatial effects observed in Fig.6(e). Furthermore, emerging methods^36,37^ suggest that it may also be possible to estimate axon diameter index from multi-TE dMRI data, which could be more sensitive to smaller axons compared to demonstration of multi-diffusion time data here.

## Discussion

We present the most comprehensive dMRI datasets collected in *ex vivo* primate brains to date, spanning whole human hemispheres and macaque brains with multi-resolution and multi-dimensional acquisitions, leveraging extended post-mortem imaging on ultra-high-gradient systems. By jointly optimizing spatial resolution, diffusion encoding, and SNR, we obtain measurements that are sensitive to fine-scale features, such as intralaminar cortical structure, hippocampal subfields, and densely packed projection pathways, that have been difficult to resolve in previous whole-brain dMRI studies. The multidimensional diffusion encoding further enhances sensitivity to microstructural differences, enabling richer modeling of tissue composition and geometry.

This represents a technological advance that is broadly valuable across scientific communities. **(i) Comparative neuroanatomy:** Whole-brain imaging in macaque and human specimens facilitates detailed cross-species comparisons and investigations of neuroanatomy, including long-range projections such as cortico-subcortical tracts and short-range, superficial fiber bundles. **(ii) Translational neuroscience:** These high-resolution, multi-dimensional data can support construction of multi-modal brain atlases, which can be used to align both gray and white-matter structures across subjects, and therefore enhance population and clinical studies. **(iii) Methodological development:** Dense sampling across spatial and diffusion-encoding dimensions provides an unprecedented testbed for the development of advanced microstructural modeling, super-resolution reconstruction, and mesoscale tractography algorithms. **(iv) Model validation:** As part of the BRAIN CONNECTS Center for Large-Scale Imaging of Neural Circuits (LINC), the same specimens will be imaged with multiple microscopic modalities. The macaque brains have received tracer injections prior to dMRI and will be imaged with polarization-sensitive optical coherence tomography (PS-OCT) and lightsheet fluorescence microscopy (LSFM). The human hemispheres will be imaged with light microscopy (PS-OCT and LSFM) or X-ray microscopy (hierarchical phase-contrast tomography and micro-CT). These datasets can support rigorous cross-modal validation and, ultimately, improvement of non-invasive dMRI methods.

While prior, open-access dMRI datasets exist for marmoset^12^, macaque^15^, chimpanzee^13^, and human brains^14^, our dataset is unique in combining ultra-high resolution, ultra-high b-values, and multidimensional sampling, in both macaque and human. Achieving these acquisition parameters in whole hemispheres or brains would not have been feasible without ultra-high-gradient MRI systems. Only one previously reported whole human brain dMRI dataset has higher nominal spatial resolution (0.2 mm isotropic)^11^. This was achieved by sectioning the brain into small blocks and scanning the blocks over multiple years on an 11.7T small-bore scanner, at substantially lower diffusion weightings. While it might have been possible to further increase the spatial resolution of our datasets by limiting the acquisition to more modest b-values, we opted for the current trade-off to allow a broader gamut of microstructural analyses.

Despite the unprecedented detail of this dataset, some limitations remain. Layer I of the cortex remains difficult to visualize directly due to its thinness (<0.4 mm). Estimates of microstructural parameters such as axon diameter index from dMRI remain challenging and SNR- and model-dependent. Post-mortem fixation also alters tissue properties, and in particular diffusivity and T2 values. More work is needed to clarify these effects and better understand the extent to which our findings extend to *in vivo* dMRI. This will be a crucial step before the multimodal data collected here can be used, *e.g.,* to train machine learning models for improved inference of brain microstructure from *in vivo* dMRI. Furthermore, the current dataset focuses on a small number of specimens and may not be appropriate for assessing individual variability. However, it provides a framework that can be scaled to larger sample sizes in the future.

These novel, multi-dimensional dMRI data of unprecedented resolution and microstructural sensitivity can drive methodological development, cross-species neuroanatomical research, and translational neuroscience. The combined potential for high-resolution tractography, microstructure mapping, and multimodal validation in the same brains opens new avenues for linking cellular architecture to whole-brain structural networks and enabling new discoveries in non-human and human primate neuroanatomy.

## Methods

### Specimen preparation

#### Human hemispheres

We included two human hemispheres in this study. One right hemisphere (Ha1) was obtained from a 70-year-old male donor, with postmortem interval of 12 hours, and one left hemisphere (Hb1) was obtained from a 63-year-old male donor, with postmortem interval of 7 hours. Tissue was obtained through approved donation protocols, and all procedures involving human tissue complied with institutional guidelines and relevant ethical regulations at Massachusetts General Hospital.

The hemispheres were fixed in 10% formalin for at least two months before MRI acquisition. Before scanning, we sealed each hemisphere in a plastic bag filled with *periodate-lysine-paraformaldehyde (PLP).* We chose PLP to ensure compatibility with downstream X-ray imaging. While it introduced background signal in b=0 images, this signal was effectively suppressed by diffusion-weighting gradients.

#### Macaque brains

We included brains from four adult male macaques (M1-M4; Macaca fascicularis). Animals were deeply anesthetized and perfused with saline followed by a fixative solution containing 4% paraformaldehyde and 1.5% sucrose in 0.1M phosphate buffer (pH 7.4). Brains were postfixed overnight and cryoprotected in increasing gradients of sucrose (10%, 20%, and 30%).

Surgery and tissue preparation of macaque brain samples were performed at the University of Rochester in accordance with the Institute of Laboratory Animal Resources Guide for the Care and Use of Laboratory Animals and approved by the University of Rochester Committee on Animal Resources. Details of the procedures are described in previous studies^38,39^.

During MRI acquisition, the macaque brain samples were packed in Fomblin (Solvay, Italy) to eliminate background signal and susceptibility artifacts.

### Imaging protocols

This work builds on our prior experience with *ex vivo* dMRI protocol optimization using high-gradient preclinical systems^20,38,40^. For the human hemispheres, we translated these protocols to the Connectome 2.0 3T MRI scanner (MAGNETOM Connectom.X, Siemens Healthineers, Forchheim, Germany), equipped with G_max_=500 mT/m and maximum slew rate=600 T/m/s, and adapted them to the larger sample size. For the macaque brains, we improved upon our previously used protocols on a Bruker 4.7 T preclinical system with G_max_=660 mT/m, in terms of both image resolution and multidimensional sampling. We used a custom-built, 64-channel *ex vivo* coil with integrated temperature stabilization for extended acquisitions on the Connectome 2.0^19^, and a product 4-channel coil on the preclinical system. On both systems, we used a 3D EPI sequence with high numbers of segments to achieve the shortest TE possible within the available scan time, and we used the maximum gradient strength of each system to reach the highest possible b-value with the shortest possible diffusion time.

### Human dMRI data acquisition

#### High-resolution protocol

We acquired high-resolution dMRI data at 0.4 mm isotropic resolution using 16 segments (TE = 43 ms; TR = 500 ms). We collected diffusion-weighted images with 64 gradient directions at b = 4,000, 8,000, and 12,000 s/mm², and 128 directions at b = 25,000 s/mm². The diffusion gradient pulse width and separation were δ = 10.4 ms and Δ = 19.5 ms, respectively. We interleaved 14 b = 0 images throughout the acquisition and acquired one additional b = 0 image with reversed phase-encoding direction for distortion correction. Total acquisition time per hemisphere was approximately 130 hours. Furthermore, to avoid heat accumulation in the RF coil during long scans at the highest b-values, we split the acquisition into sessions with interleaved breaks and distributed the scans with the highest b-values across these sessions. Thus, data acquisition with this protocol took a week to complete.

#### Multi-diffusion-time protocol

We acquired multi-shell data at two diffusion times for microstructural modeling using 8 segments and 0.8 mm isotropic resolution (TE = 49 ms; TR = 500 ms). For the shorter diffusion time (δ/Δ = 11.5/17.5 ms), we collected 32 directions at b = 2,000, 4,600, 8,100, 12,700, and 18,300 s/mm², and 64 directions at b = 25,000 s/mm². For the longer diffusion time (δ/Δ = 11.5/25.5 ms), we additionally acquired 64 directions at b = 31,000 and 41,000 s/mm². We interleaved b=0 images every 32 directions and acquired one additional b=0 image with reversed phase-encoding direction for distortion correction. Total acquisition time was approximately 63 hours.

#### Multi-echo-time protocol

We acquired multi-shell data at four echo times to support combined diffusion–relaxometry analysis. Using 6 segments, we acquired 1.0 mm isotropic resolution data with TEs of 42, 55, 70, and 88 ms (TR = 500 ms). For each TE, we collected 32 directions at b = 2,000, 4,600, 8,100, 12,700, and 18,300 s/mm², and 64 directions at b = 25,000 s/mm². We interleaved b=0 images every 32 directions and acquired one additional b=0 image with reversed phase-encoding direction at each of the four TEs for distortion correction. Total acquisition time was approximately 56 hours.

### Macaque dMRI data acquisition

#### High-resolution protocol

We acquired high-resolution macaque dMRI data at 0.25 mm isotropic resolution using 16–18 segments (TE ∼ 28 ms; TR = 500 ms; δ/Δ = 9/13 ms), with maximum b-values of 20,000 s/mm². We implemented two q-space sampling schemes:

i. **Multi-shell sampling.** We acquired four shells at b = 4,000, 8,000, 12,000, and 20,000 s/mm² using 32, 32, 64, and 64 directions, respectively. We interleaved b=0 images every 32 directions. Total acquisition time was approximately 78 hours.
ii. **Under-sampled Cartesian grid scheme.** We acquired 171 diffusion directions, sampled randomly from a 11^3^ Cartesian lattice enclosed in a sphere and one b=0 image. These can be used to reconstruct fully sampled, 514-direction diffusion spectrum imaging data, e.g., using compressed sensing^41^. Total acquisition time was approximately 60 hours.

Acquiring the two schemes in the same brains provides a resource for future studies to evaluate data sampling strategies, e.g., for fast acquisition, and fiber orientation reconstruction methods. We acquired data with the under-sampled grid scheme from all four macaque specimens, and the multi-shell scheme from three (M2-M4).

#### Multi-echo/multi-diffusion time protocol

We acquired data at 0.5 mm isotropic resolution using 8 segments (TR = 500 ms), sampling three diffusion times and multiple echo times. For the shortest diffusion time (δ/Δ = 9.6/12 ms), we acquired seven shells at b = 1000, 2500, 5000, 7500, 11100, 18100, and 25000 s/mm² and TE = 28 ms. For the intermediate diffusion time (δ/Δ = 11/15 ms), we acquired eight shells, including the seven above and one at b = 43,000 s/mm², at TEs of 33, 45, 60, and 78 ms. For the longest diffusion time (δ/Δ = 11/21 ms), we acquired nine shells, including the seven above and two at b = 43,000 and 64,000 s/mm², at a TE of 38 ms. Total acquisition time was approximately 90 hours.

### T2 mapping data acquisition

We acquired spin-echo images at multiple TEs to estimate the T2 constant. This was useful both for guiding the optimization of the dMRI protocols with respect to gray-white matter contrast and SNR, and for allowing more precise modeling of microstructural tissue parameters.

For human specimens, we acquired spin-echo multi-contrast (SEMC) images at 1.1 mm isotropic resolution, with 2D encoding, no partial Fourier, whole-hemisphere coverage, 10 concatenations. The spin-echo images were collected at 32 echo times with an echo spacing of 7.6 ms and TR of 3000 ms. Total image acquisition time was approximately 1 hour.

For macaque specimens, we acquired multi-slice multi-echo (MSME) images with slice selective RF pulses and 3D encoding at 0.5 mm isotropic resolution, without partial Fourier or signal averaging. The spin-echo images were collected at 32 echo times with an echo spacing of 6 ms and TR of 500 ms. Total imaging time was approximately 1.5 hours.

### dMRI data preprocessing

We preprocessed all dMRI data using a minimal standardized pipeline adapted to *ex vivo* human and macaque acquisitions. For all MRI datasets, we reoriented volumes to the correct anatomical orientation.

#### Human data

We denoised the human dMRI data^42^ and corrected for distortions due to gradient nonlinearity^43^, B0 field inhomogeneity^44^, eddy currents^45^, B1 field inhomogeneity^46^, and signal drift^47^. To correct signal drift, we evaluated both linear and quadratic drift models and selected the model that maximized temporal signal-to-noise ratio (tSNR) in the interleaved b = 0 images.

#### Macaque data

Similarly to the human dMRI data, we performed denoising and correction for distortions due to eddy currents and B1 field inhomogeneity. Signal drift was negligible on the preclinical system and therefore not corrected.

For multi-dimensional dMRI acquisitions (multiple echo times and/or diffusion times), we preprocessed data collected with each echo or diffusion time independently. Before combining the data, we checked the position of the first b=0 reference volume for each echo or diffusion time and applied affine registration when necessary. In practice, positional changes between sessions were negligible in the human datasets, and we did not perform such registration.

When combining human data acquired at multiple diffusion times and same TE, we verified signal consistency across scans using b = 0 images and applied additional signal drift correction if necessary.

### Human image segmentation

#### Brain segmentation

We segmented cortical and subcortical brain structures on the 0.4 mm isotropic dMRI data automatically, using the average of b=4000 s/mm^2^ images normalized by the average of the b=0 s/mm^2^ images, which has T1-like contrast. We employed segmentation tools available in FreeSurfer, including the classical recon-all pipeline^48^ and SAMSEG with the NextBrain atlas^49^. An existing parcellation^50^ of the prefrontal cortex based on cytoarchitectonic features^51^ was mapped from the ‘fsaverage’ template and to the individual surface in dMRI space. We used this parcellation to segregate tractography streamlines of the internal capsule based on their cortical termination.

#### Hippocampal subfield annotation

As the automated whole-brain segmentations did not delineate the boundaries of hippocampal subfields and substructures with satisfactory accuracy, we annotated them manually on the normalized mean b=4000 s/mm^2^ images at 0.4 mm isotropic resolution. We followed the pentad segmentation protocol^27^ and additional histological resources^52,53^ to label hippocampal regions. The image contrast allowed us to parcellate the CA1-CA4, subiculum hippocampal-amygdaloid transition area (HATA), subiculum, fimbria, alveus, granular layer of the dentate gyrus and the stratum radiatum and stratum lacunosum-moleculare (SRLM). The dentate gyrus appeared darker, hippocampal subfields exhibited intermediate intensity, and SRLM appeared brighter, whereas the alveus and fimbria appeared brightest. Along the allocortical ribbon, boundaries between subfields of CA1-CA4 and HATA were approximated based on typical locations along the anterior-posterior axis^27^. We grouped the subicular subfields (subiculum, prosubiculum, presubiculum and parasubiculum) into one subiculum label.

#### Cortical layer segmentation

We segmented the upper (supragranular) and lower (infragranular) cortical layers using a semi-supervised deep-learning approach previously developed for high-resolution 7T structural images with T2-weighted contrast^28^. To match the required input resolution, we up-sampled our 0.4 mm isotropic mean b=0 image to 0.12 mm isotropic, after further correction for intensity bias field using ANTs^46^. We followed an overlapping-patch strategy, averaging local predictions from partially overlapping regions to ensure spatial continuity, minimize boundary artifacts, and produce detailed laminar segmentations for analysis and visualization of microstructure variations.

### Macaque image segmentation

We used the cortical and subcortical segmentations included in the CIVM macaque template^54^. We used ANTs^55^ to perform non-linear registration between each macaque’s T2-weighted MSME images collected with TE=30 ms and the CIVM T2-weighted template image, as well as between the macaque’s MSME and the average b=0 images from the high-resolution scan at 0.25 mm and multi-dimensional dMRI scan with shortest TE at 0.5 mm. We applied the resulting deformation fields to transform the CIVM atlas segmentations into the macaque high-resolution and multi-dimensional dMRI space, respectively. We used these segmentations to segregate tractography streamlines of the internal capsule based on their terminations. We also mapped the hippocampal annotations of the CIVM template^56,57^ to each macaque brain to guide annotations of hippocampal tracts.

### Tractography

We performed fiber orientation estimation and tractography analysis using MRtrix3^58^. Using a multi-shell, multi-tissue constrained spherical deconvolution (MSMT-CSD) approach^59^, we reconstructed fiber orientation distribution functions (fODFs) from the high-resolution human and macaque dMRI data. We used a binary tissue mask including only white and gray matter voxels to constrain voxel selection for estimation of tissue-specific response functions.

We then performed probabilistic tractography using fODFs, propagating streamlines from five random seed locations within each white-matter voxel to generate whole-brain tracts. For visualization, we generated direction-encoded color maps from the white-matter fODFs^22^ and tract density images from the whole-brain tracts^60^. For visualizing hippocampal fibers, we repeated tractography with denser seeding (ten seed locations per voxel) restricted to voxels in the hippocampus. We delineated specific fiber bundles in TrackVis based on regions of interest (ROIs) from the aforementioned segmentations.

### Microstructure model fitting

We performed microstructure model fitting using the Julia package Microstructure.jl^26^.

#### Soma and neurite density imaging (SANDI)

We fit the SANDI model^25^, a three-compartment model consisting of (i) a sphere compartment modeling water diffusion in cell bodies, (ii) a stick compartment modeling water diffusion in dendrite-like structures, and (iii) an isotropic tensor compartment representing diffusion in the extra-cellular space. The free parameters of the model include the intra-soma signal fraction *f_soma_*, soma radius *R_soma_*, intra-neurite signal fraction *f_neurite_*, intra-neurite parallel diffusivity 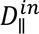 and diffusivity of the isotropic extra-cellular space *D_ec_*_’-_. Based on experimental observations in *ex vivo* tissue, we fixed 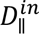 to 0.6 *μ*m^2^/ms.

*Ex vivo* tissue can exhibit a non-decaying signal component associated with fixation-related immobile water, commonly referred to as the “dot” compartment. To account for this effect, we evaluated both the standard three-compartment SANDI model and a four-compartment SANDI-dot model that includes an additional dot compartment. In our samples, the estimated signal fraction of the dot compartment was negligible (<0.05) in gray matter and approximately 0.1 in white matter. The dot compartment is thus necessary when displaying whole-brain microstructure maps. Including the dot compartment, however, increased noise in the microstructure map of the high-resolution human data due to the limited number of b-values. For datasets with more b-values, including the dot compartment did not affect estimation accuracy.

We estimated model parameters using a probabilistic neural network estimator trained on synthetic data. Leveraging prior knowledge that *ex vivo* tissue exhibits reduced extracellular water fractions and minimal dot contributions, we constrained the training distributions to improve estimation accuracy for intra-soma and intra-neurite fractions. We evaluated multiple model configurations and selected the one that produced the most reliable estimates from our four-shell acquisition, based on validation using synthetic datasets (Extended Data Fig.5).

We generated 50,000 training samples with tissue parameters uniformly sampled from prior ranges of [3, 12] *μ*m for soma radius in humans, [1, 10] *μ*m for soma radius in macaque, and [0.2, 1.2] *μ*m^2^/ms for extracellular diffusivity in both species. The compartment signal fractions were sampled from a Dirichlet distribution, with lower concentration parameters for the extra-cellular and dot compartments (0.3 v.s. 1.0 for intra-soma/-neurite compartments), which represents the prior knowledge of lower extra-cellular/dot signals in *ex vivo* tissue. We generated synthetic signals using the forward model and added Gaussian noise based on the temporal SNR of the b=0 images and the number of measurements used for direction averaging.

#### Multi-TE SANDI (MTE-SANDI)

The MTE-SANDI model^32^ extends SANDI by explicitly modeling compartment-specific T2 relaxation times. This enables estimation of unbiased soma and neurite density indices, as opposed to the T2-weighted signal fractions derived from single-TE acquisitions. In addition to microstructural indices, MTE-SANDI provides compartment-specific T2 estimates for the intra-soma, intra-neurite, and extra-cellular compartments, which reflect tissue composition. For MTE-SANDI analysis, we focused on the gray matter, where the dot compartment was negligible, as including the T2 of the dot compartment would lead to more model parameters than can be robustly differentiated based on the b-values and TEs in our data.

We again used a neural network estimator trained on synthetic data and optimized model configurations to achieve robust parameter estimation with the four-TE acquisition protocol. Evaluation and model comparisons are shown in Extended Data Fig.6.

To accommodate the increased model dimensionality, we generated 500,000 training samples and trained a larger neural network. Tissue parameters were uniformly sampled from prior ranges of [3, 12] *μ*m for soma radius in humans, [1, 10] *μ*m for soma radius in macaques, [0.2, 1.2] *μ*m^2^/ms for extracellular diffusivity, and [20, 100] ms for the T2 of all compartments in humans. For macaques, we used a narrower T2 range of [20, 70] ms, based on the T2 differences between species that we observed in our T2 mapping data. The compartment fractions were sampled from a Dirichlet distribution with lower concentration parameters for the extra-cellular compartment (0.3 v.s. 1.0 in intra-soma/intra-neurite compartments), incorporating prior knowledge of reduced extracellular water in *ex vivo* tissue. We added Gaussian noise to the simulated signals based on the temporal SNR of the b=0 images and the number of measurements used for direction averaging at the shortest TE.

#### Axon diameter index

We estimated the axon diameter index using a three-compartment model that we recently evaluated for *ex vivo* tissue^20,26^. The model consists of (i) a cylindrical compartment, modeling restricted water diffusion within axons, (ii) a tensor compartment, modeling hindered water diffusion in extra-cellular space, and (iii) a dot compartment, accounting for immobile water in the white matter. We estimated model parameters using a Markov chain Monte Carlo (MCMC) sampler. Detailed descriptions of the model and estimation framework are provided in our prior work, which demonstrates that axon diameter index estimation is highly sensitive to SNR. Synthetic evaluations show that both accuracy and precision improve with high SNR and multiple diffusion-time measurements. We demonstrated the estimation of axon diameter index using the human two-diffusion time data in comparison to using only the longer diffusion time data.

## Data Availability

The diffusion MRI data and derived maps in this study will be openly available on the DANDI Archive (RRID:SCR_017571). The human data will be available at https://dandiarchive.org/dandiset/001278 and the macaque data will be available at https://dandiarchive.org/dandiset/001372.

## Code Availability

Code that generates key results of the study will be available at https://github.com/lincbrain. We used publicly available software tools for the data analysis, including FSL (https://fsl.fmrib.ox.ac.uk/fsl), MRtrix3 (https://www.mrtrix.org), Microstructure.jl (https://github.com/Tinggong/Microstructure.jl) and FreeSurfer (https://github.com/freesurfer/freesurfer).

## Acknowledgement

This work is supported by the center for Large-scale Imaging of Neural Circuits (LINC), an NIH BRAIN Initiative Connectivity across Scales (CONNECTS) comprehensive center (UM1-NS132358). Additional support is provided by the NIH National Institute of Biomedical Imaging and Bioengineering (K99-EB037396) and National Institute of Neurological Disorders and Stroke (R01-NS119911, R01-NS127353).

## Author contribution

TG contributed to the development of the imaging protocols, acquisition of human and macaque MRI data, conceptualization and administration of the study, analysis and visualization of the data, and writing of the manuscript; CM contributed to tractography analysis and visualization; DS contributed to acquisition of human MRI data; EB, JW, JS contributed to acquisition and preprocessing of macaque MRI data; EB and ER contributed to annotation of hippocampus subfields; XZ contributed to segmentation of cortical layers; GR, AM, MM, BK contributed to development and testing of the human ex vivo coil; KG and SG contributed to the standardization and distribution of MRI data; JCA contributed to human specimens and provided anatomical guidance; SYH contributed to human MRI resources; SNH contributed to macaque specimens and funding acquisition; AY contributed to conceptualization and administration of the study, editing of the manuscript, and funding acquisition. All authors provided revisions and approved the manuscript for publication.

