## Extended Data for "Multi-dimensional diffusion MRI at ultra-high gradient strength for mapping axonal architecture and microstructure in the primate brain"

### A. Human dMRI protocols on Siemens Connectome 2.0 3.0T system ($G_{max} = 500\ mT/m$)

| Protocol | Resolution (mm³) | Segments | TR (ms) | TE (ms) | δ / Δ (ms) | b-values (s/mm²) | # Directions per b-shell | Special features | Acquisition Time |
|---|---|---|---|---|---|---|---|---|---|
| **High-resolution** | 0.4 | 16 | 500 | 43 | 10.4 / 19.5 | 0, 4000, 8000, 12000, 25000 | 64 (low b), 128 (for b = 25000) | Multi-shell with dense orientation sampling | ~130 h |
| **Multi-diffusion-time** | 0.8 | 8 | 500 | 49 | 11.5 / 17.5 (S1)<br>11.5 / 25.5 (S2) | 0, 2000, 4600, 8100, 12700, 18300, 25000, 31000, 41000 | 32 (low b), 64 ( for b = 25000, 31000, 41000) | Maximum b-value for shorter diffusion time is 25K; | ~63 h |
| **Multi-echo-time** | 1.0 | 6 | 500 | 42 (S3)<br>55 (S4)<br>70 (S5)<br>88 (S6) | 11.5 / 17.5 | 0, 2000, 4600, 8100, 12700, 18300, 25000 | 32 (low b), 64 (for b = 25000) | Diffusion sampling repeated for four TEs | ~56 h |

### B. Macaque dMRI protocols on Bruker 4.7T system ($G_{max} = 660\ mT/m$)

| Protocol | Resolution (mm³) | Segments | TR (ms) | TE (ms) | δ / Δ (ms) | bmax (s/mm²) | # Directions | Special features | Acquisition time |
|---|---|---|---|---|---|---|---|---|---|
| **High-resolution** | 0.25 | 17 | 500 | 28 | 9.0 / 13.0 | 20000 | 32 (b=4k, 8k)<br>64 (b=12k, 20k) | Multi-shell with dense orientation sampling | ~ 78 h |
| **High-resolution** | 0.25 | ~16 | 500 | 28 | 9.0 / 13.0 | 20000 | 171 (on under-sampled Cartesian grid) | Diffusion Spectrum imaging data can be recovered | ~60 h |
| **Multi-echo / Multi-diffusion-time** | 0.5 | 8 | 500 | 28 (S1)<br>33 (S2)<br>38 (S3)<br>45 (S4)<br>60 (S5)<br>78 (S6) | 9.6 / 12.0<br>11.0 / 15.2<br>11.0 / 21.0<br>11.0 / 15.2<br>11.0 / 15.2<br>11.0 / 15.2 | 25000<br>43000<br>64000<br>43000<br>43000<br>43000 | 12 (for b = 1000, 2500, 5000, 7500)<br>32 (for b = 11100, 18100, 25000, 43000, 64000) | Diffusion sampling repeated for 3 diffusion times and 4 TEs | ~86 h |

### C. T2 distributions in macaque and human brain

| Tissue T2 | WM | GM |
|---|---|---|
| **Human** | ~58 ms | ~72 ms |
| **Macaque** | ~37 ms | ~42 ms |

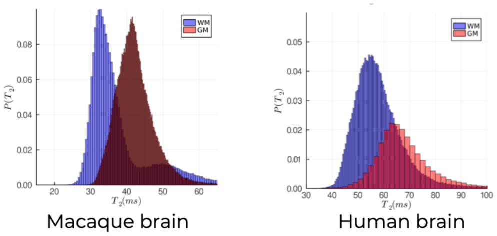


Extended Data Fig.1 | Summary of dMRI protocols. (A) human protocols; (B) macaque protocols (C) T2 distributions in the white-matter (WM) and gray-matter (GM) of a typical macaque and human brain sample.

Extended Data Table 1 | Mean temporal SNR of preprocessed dMRI data from two human samples. S1–S6 sequentially represent the SNR for each diffusion-time session in the two-diffusion-time scan and each echo-time session in the four-echo-time scan, as shown in Extended Data Fig. 1A.

| | **High-resolution data** | **Multi-dimensional data** | | | | | |
|---|---|---|---|---|---|---|---|
| | | **S1** | **S2** | **S3** | **S4** | **S5** | **S6** |
| Ha1 | 35.2 | 39.4 | 30.8 | 61.6 | 65.8 | 54.4 | 46.6 |
| Hb1 | 23.4 | 60.9 | 45.4 | 68.5 | 46.1 | 43.6 | 32.8 |

Extended Data Table 2 | Mean temporal SNR of preprocessed dMRI data from four macaque samples (M1-4). The SNR values for the high-resolution data were calculated from multi-shell scans containing multiple b0 images. S1–S6 sequentially represent the SNR for each session in the multi-echo, multi-diffusion-time scan shown in Extended Data Fig. 1B.

| | **High-resolution data** | **Multi-dimensional data** | | | | | |
|---|---|---|---|---|---|---|---|
| | | **S1** | **S2** | **S3** | **S4** | **S5** | **S6** |
| M1 | | 51.3 | 74.6 | 62.0 | 46.7 | 34.0 | 29.5 |
| M2 | 41.7 | 33.7 | 75.3 | 68.4 | 59.0 | 39.6 | 29.9 |
| M3 | 50.7 | 73.4 | 78.7 | 76.6 | 65.4 | 47.9 | 33.6 |
| M4 | 42.9 | 41.2 | 84.6 | 69.7 | 56.3 | 44.2 | 32.0 |
| mean | 45.1 | 49.9 | 78.3 | 69.2 | 56.9 | 41.4 | 31.3 |

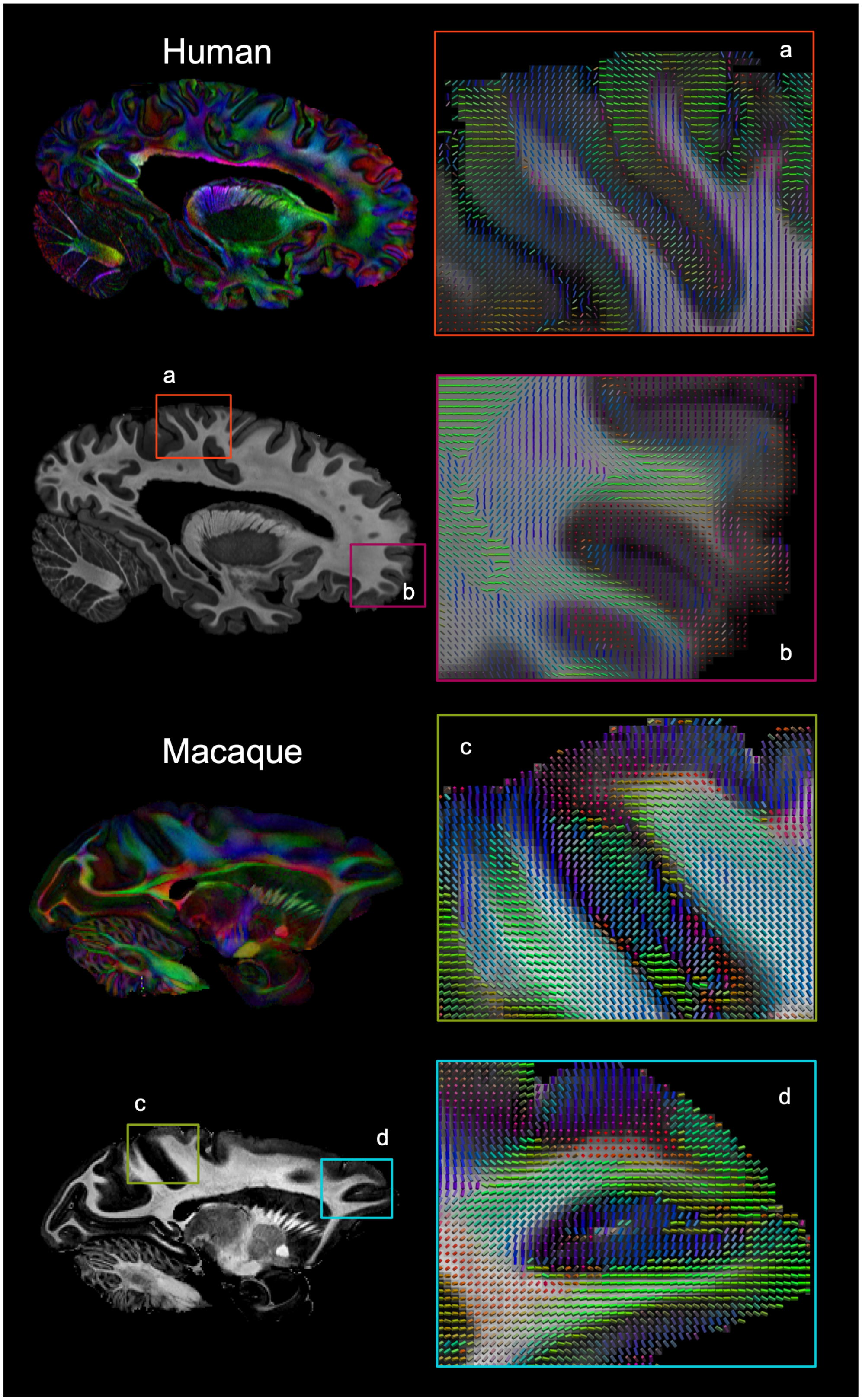


Extended Data Fig. 2 | Principal fiber orientations from diffusion tensor imaging at gray-white matter interface.

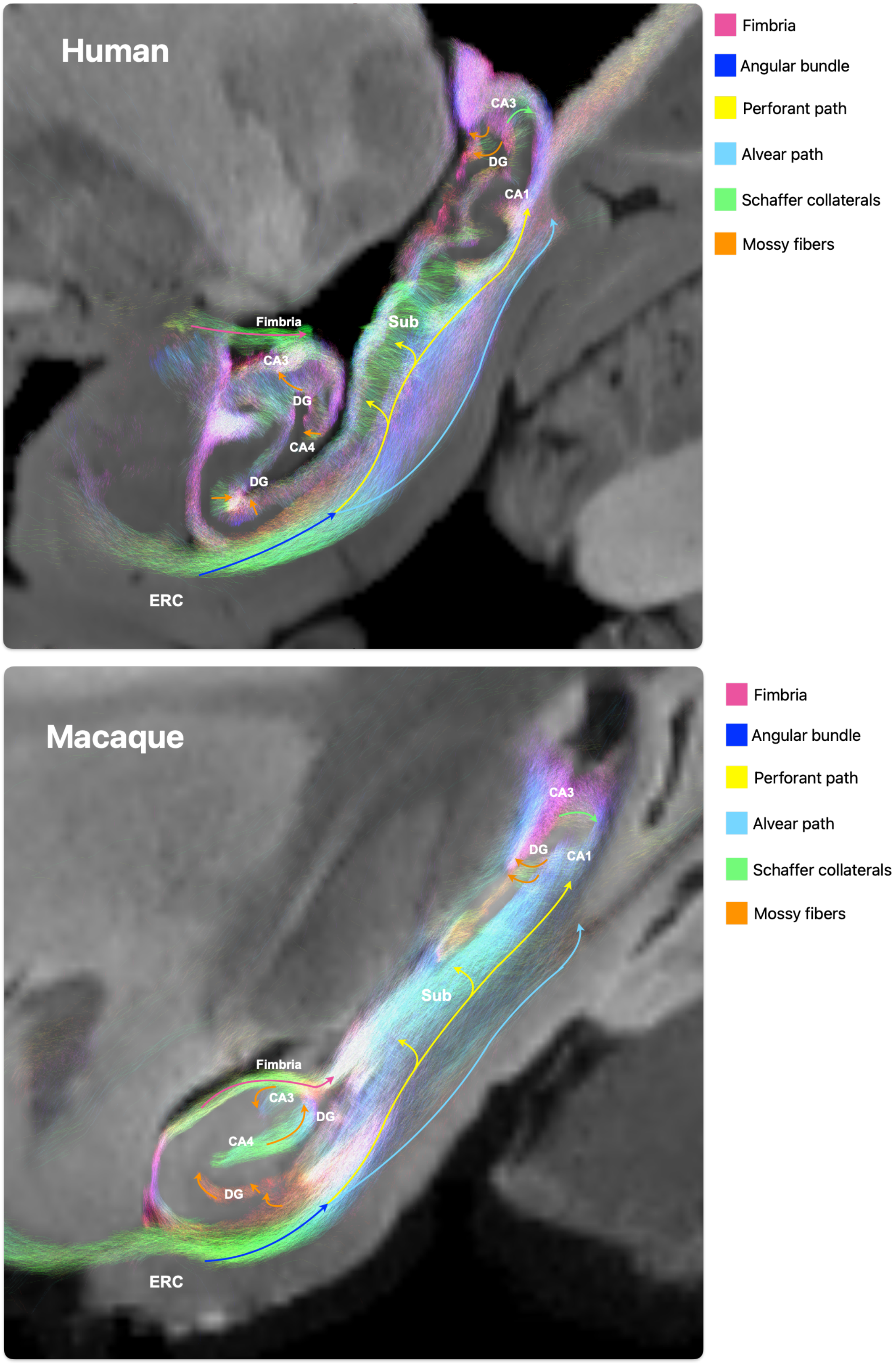


Extended Data Fig.3 | Full depiction of hippocampal tracts. Macaque cortex has less gyrification compared to human.

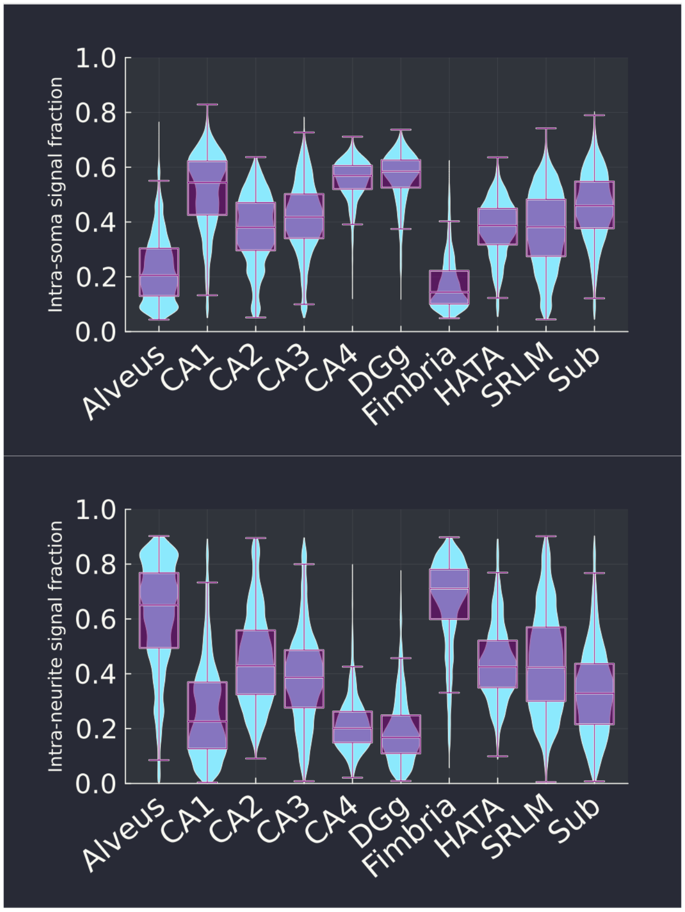


Extended Data Fig.4 | Distributions of intra-soma and intra-neurite signal fractions in the hippocampal substructures from human sample Hb1.

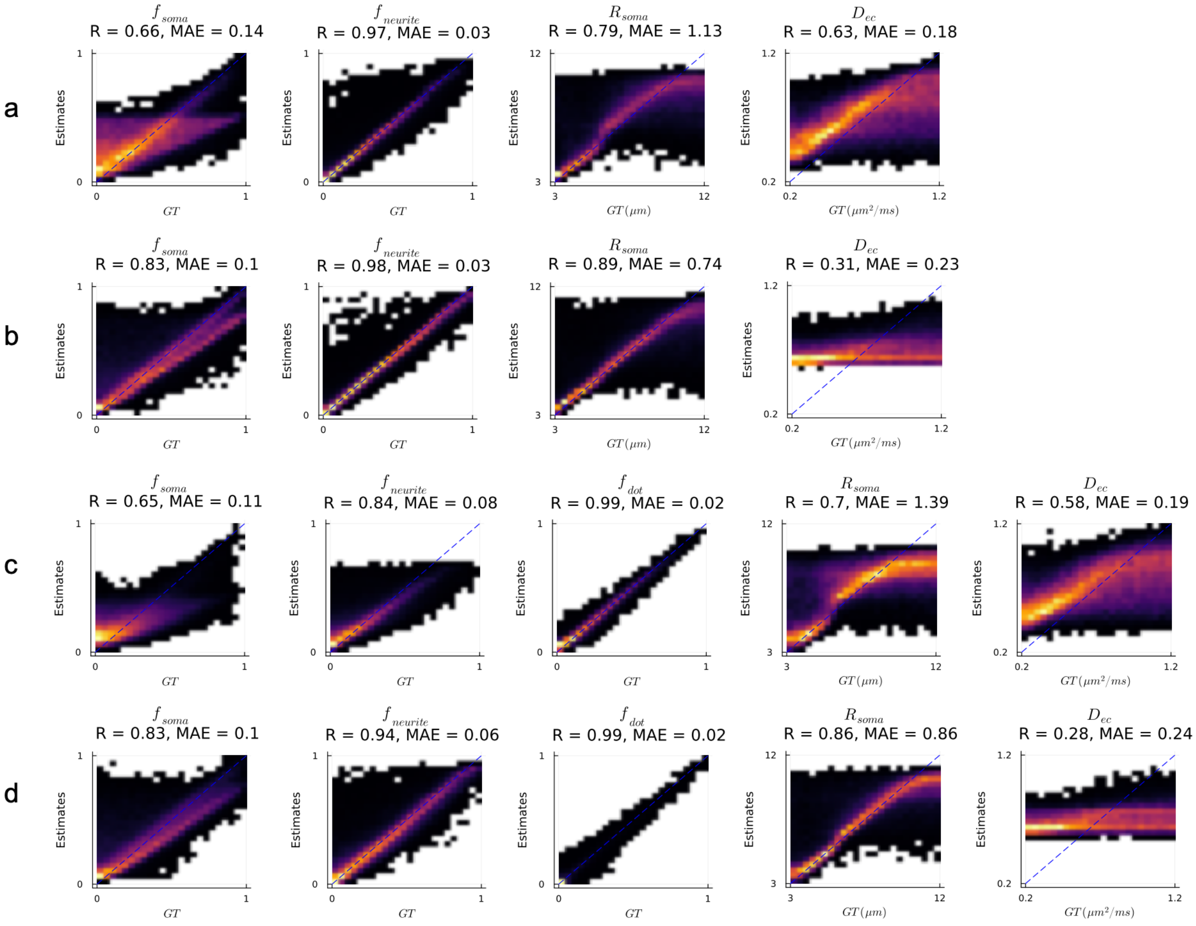


Extended Data Fig.5 | Optimization for intra-soma and intra-neurite signal fraction estimation on human high-resolution protocol. Evaluation was performed on synthetic data, where ground truth (GT) values v.s. estimates were displayed as heatmaps along with their correlation coefficient (R) and mean absolution error (MAE). Simulations were performed for the SANDI model trained on (a) uniform concentration across compartments and (b) lower concentration for the extra-cellular compartment; and the SANDIdot model trained on (c) uniform concentration and (d) lower concentration for the extra-cellular and dot signal compartments.

A lower concentration means that fewer samples with high extra-cellular or dot signal fractions are included as training data. For both SANDI and SANDIdot, incorporating prior information of low extra-cellular and dot signal fractions improved the estimation of the intra-soma/intra-neurite signal fraction and soma radius. Though the low concentration of the extra-cellular signal also made extra-cellular diffusivity less reliable, it is not the parameter of interest.

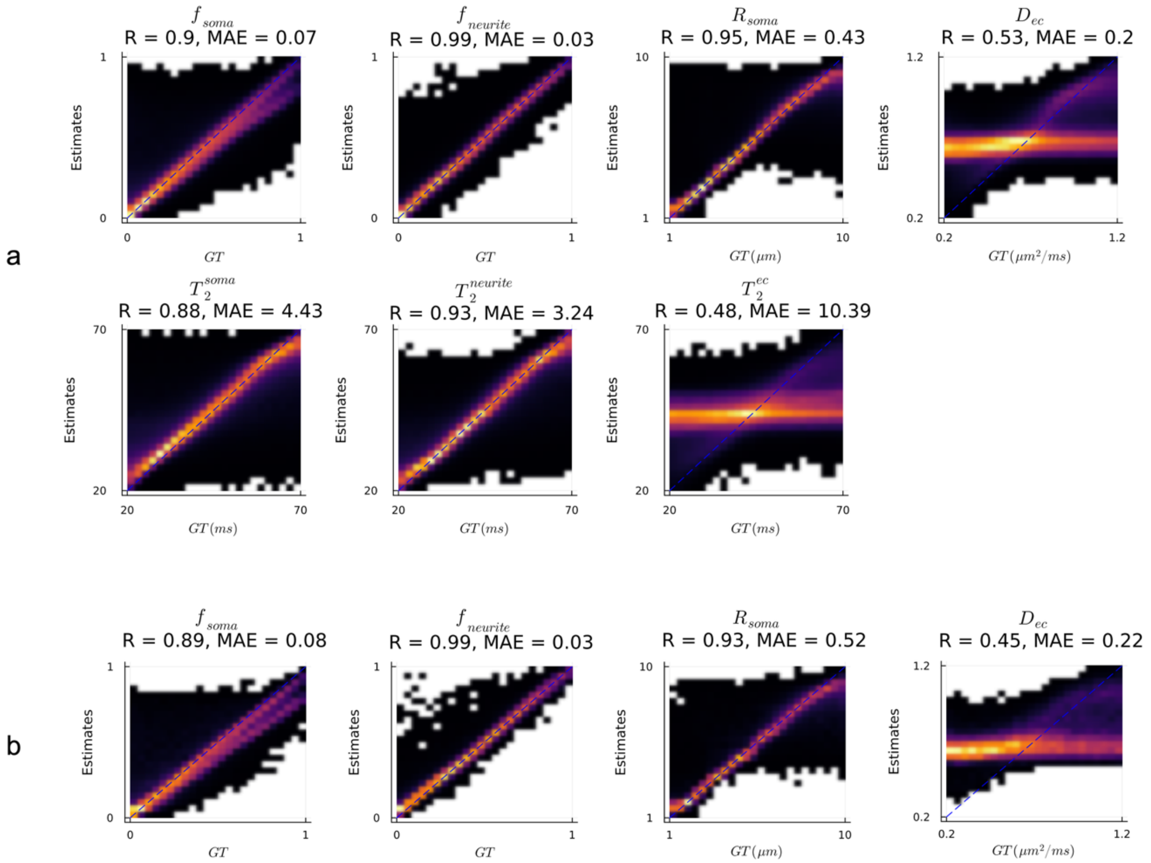


Extended Data Fig.6 | MTE-SANDI fitting evaluations on macaque data protocol with four different TEs (a) in comparison to using only data from the shortest TE (b). Similarly to the previous evaluation, we used a lower concentration for the extra-cellular compartment to optimize accuracy in the estimation of the parameters associated with the intra-soma and intra-neurite compartment. Compared to single-TE SANDI, MTE-SANDI improved parameter accuracy and precision for the intra-soma signal fraction and soma radius.
